# Cultivation and molecular characterization of foley catheterized urine, clean catch urine, and vaginal swabs from pregnant women prior to delivery

**DOI:** 10.1101/2024.12.05.627013

**Authors:** Jonathan M. Greenberg, Roberto Romero, Ali Alhousseini, Andrew D. Winters, Adi L. Tarca, Nicole M. Gilbert, Nardhy Gomez-Lopez, Sonia S. Hassan, Kevin R. Theis

## Abstract

The bladder and urine have historically been considered sterile, especially in the context of clinical assessment. Yet, enhanced culture techniques and advances in DNA sequencing surveys have revealed a low microbial biomass microbiota in the bladders of most healthy individuals. Yet very few studies have investigated the urinary microbiota of pregnant women, who are at increased risk of urinary tract infections (UTIs) that can lead to pregnancy complications such as spontaneous preterm birth. To better understand the potential role of a bladder microbiota during pregnancy, we characterized the urinary microbiota of 25 pregnant women (delivering at or beyond 36 weeks gestation) by comparing the bacterial profiles of their paired catheter urine, clean catch urine, and vaginal swabs through both cultivation and molecular microbiological survey methods. For culture, three bacterial taxa were detected in at least 20% of all urine samples (*Lactobacillus* species, coagulase negative *Staphylococcus* species, and *Ureaplasma urealyticum*). All three taxa were detected less frequently in Foley catheter urine than in clean catch urine. *Ureaplasma urealyticum* was the most frequently recovered bacterium in Foley catheter urine (13/25 women). It was also identified as being more relatively abundant in Foley catheter urine than in the vagina through 16S rRNA gene surveys. Other lower abundance Gram-positive anaerobic cocci (i.e., *Finegoldia* and *Anaerococcus)* were also more relatively abundant in Foley catheter urine than clean catch urine or vaginal swabs. However, all sample types had high relative abundances of *Lactobacillus* and *Gardnerella* species. Overall, this suggests that, although vaginal microbiota contamination cannot be completely avoided, Foley catheterized urine is effective at characterizing the low abundance bladder microbiota, including *Ureaplasma*, *Finegoldia,* and *Anaerococcus* species. This warrants their further consideration as commensal members of the bladder microbiome during pregnancy.

## INTRODUCTION

The bladder and urine have historically been regarded as sterile, however, over the past decade, studies have demonstrated that when cultivation conditions that accommodate the atmospheric and metabolic requirements of potential resident microorganisms are provided, these microorganisms can be cultured from urine in healthy patients [1–8]. Complementing these culture-based observations, contemporary sequencing technologies, such as 16S rRNA gene and metagenomic sequencing, have afforded characterization of a potential urinary microbiota in healthy female and male subjects [6, 9–14]. Collectively, these studies provide evidence for the existence of a urinary microbiota, which may competitively and/or directly exclude potential pathogens from the urinary tract. Dysbiosis (i.e., a change or disruption of resident microbial communities) of the bladder microbiota could itself contribute to urinary incontinence, and other urinary tract disorders, or the loss of potentially protective urinary microbiota members could lead to an overgrowth of opportunistic pathogens that cause urinary tract infections (UTIs) [11]. However, whether and how the urinary microbiota contributes to disease risk necessitates further characterization of the urinary microbiota under differing physiological contexts. The composition of the urinary microbiota, and its relationship to the vaginal microbiota, in pregnant women, who are particularly at high risk for UTIs, is a notable gap in knowledge for the field [15].

Pregnancy involves a series of physiological and morphological changes in the urinary tract, including dilatation of the ureters and renal pelvis, increase in bladder pressure, higher glomerular filtration rate, and alteration of osmolality [16, 17]. Such changes lead to a greater likelihood of urinary stasis and vesicoureteral reflux [18, 19], conditions that can facilitate microbial growth [20], and thus increase the risk of UTIs. In fact, UTIs are the most common bacterial infection in women during pregnancy, occurring in up to 8% of pregnancies [21]. UTIs in pregnancy are associated with maternal and perinatal complications, including low birth weight, disturbance of the immune system, spontaneous preterm birth, and maternal sepsis [22–34]. Clinically, the presence of bacteria in the urinary tract is considered pathologic and can be classified as 1) asymptomatic bacteriuria (ASB), which is typically diagnosed by the presence of 100,000 colony forming units per milliliter of urine (CFU/ml) without any associated symptoms [35–37]), 2) UTI, which is typically diagnosed by the presence of 1,000 or 10,000 CFU/ml of urine with associated symptoms such as dysuria, urgency, or hematuria [36, 37], or 3) pyelonephritis, which is diagnosed by the presence of bacteriuria accompanied by systemic signs due to a kidney infection, such as lower back pain, fever, nausea, and urinary urgency [35–37]. While pregnant women diagnosed with ASB or UTI are under comparable risk for adverse pregnancy outcomes, the main difference between these two diagnoses, aside from symptom presentation, is the magnitude of culturable bacteria in urine, which is still the clinical diagnostic standard along with urinalysis [38]. Notably, ASB has been reported in up to 10% of pregnancies [39–43], and, if left untreated, can lead to symptomatic UTI, including pyelonephritis in 30-36% of the cases [44, 45]. Indeed, contrary to the management of healthy non-pregnant individuals [38], there is a consensus for the benefits of screening and antibiotic treatment of ASB during pregnancy to reduce the incidence of pyelonephritis and its adverse consequences [35, 39]. Yet, recent studies have indicated the presence of a urinary microbiota during uncomplicated pregnancies [46–49]. It is possible the urinary microbiota plays a physiological role in healthy pregnancy, which could be disrupted by antibiotic treatment for ASB. However, further study of the composition and load of the urinary microbiota during pregnancy is required before its potential relevance to current clinical practice can be fully appreciated.

Given that the urinary tract was thought to be sterile and that numerous recent studies [2, 14, 47, 50–53] have found evidence for a urinary microbiota, two important caveats must be recognized when characterizing the urinary microbiota of women, especially during pregnancy. First, urine samples are susceptible to vulvovaginal contamination due to the close proximity of the urethral opening to the vagina, so it is not possible to conclude that all characterized microorganisms actually reside in the bladder or the urine. To best decipher the biological source of bacteria detected in urine, is critical to include paired samples from adjacent mucosal sites such as periurethral and vulvovaginal swabs. It is also important that collected urine specimens be promptly frozen to mitigate growth and replication of bacteria acquired in urine from adjacent sites. Second, resident urinary microbial communities are likely present in very low abundances, therefore a sufficient volume of urine is needed for effective DNA extraction. Additionally, when attempting to characterize the urinary microbiota through contemporary DNA sequencing technologies, there is risk of amplifying and characterizing background bacterial DNA contamination from DNA extraction kits and PCR reagents, i.e. the “kitome” [54–58]. To properly account for these kitome signals, appropriate background technical controls need to be included in such studies.

Herein, we aimed to assess two methods of urine collection for characterization of the urinary microbiota during pregnancy with appropriate consideration being given to potential contaminating sources of bacteria and/or their DNA. Specific secondary objectives were to: 1) assess the bacterial load of urine sampled using catheter and clean catch collection methods via quantitative real-time PCR; 2) compare the composition and structure of the bacterial profiles of urine from pregnant women obtained using catheter and clean catch collection approaches with those of background technical controls; 3) assess the similarity of culture and contemporary DNA sequencing characterizations of urinary bacterial profiles; and 4) contrast the bacterial profiles of the urine of pregnant women obtained through these two collection approaches with those of the vagina to assess potential vulvovaginal contamination.

## RESULTS

### Study Component 1: Comparing the bacterial load and 16S rRNA gene profiles of different volumes of urine from the same women

#### Patient characteristics

Eight pregnant women were included in Component 1 of the study. Foley catheter urine (n = 4) was used from four women, while clean catch urine (n = 5) was used from five women. Sufficient urine volume from both sample types was collected from one woman. The median and interquartile ranges for age, body mass index (BMI), gestational age at sampling, and neonatal birthweight were 27.5 (25.2-28.2) years, 31.6 (28.5-45.8) kg/m^2^, 39.7 (38.8-40.8) weeks, and 3,392 (3,256-3,882) grams, respectively **(Supplementary Table 1)**.

#### Bacterial load assessed by quantitative real-time PCR (qPCR) in Foley catheter and clean catch urine samples exceeded technical controls and varied by urine volume processed

Urine samples from Foley catheter and clean catch collection methods were assessed via qPCR to evaluate whether bacterial load was greater than that observed in negative technical controls and whether urine volume would affect bacterial load. Median cycle of quantification (Cq) values for each volume of urine were significantly lower than the median Cq values for blank DNA extraction kit controls, regardless of whether the urine was collected via Foley catheter (Mann-Whitney; U = 0, p = 0.0058) or the mid-stream clean catch method (U = 0, p = 0.0081), indicating that urine consistently contains a detectable bacterial load through qPCR. Furthermore, the volume of urine processed had an effect on Cq value for Foley catheter (repeated measures ANOVA; F = 9.805, p < 0.0001) and clean catch (F = 28.01, p < 0.0001) urine samples. For both urine collection methods, a urine volume of 5.4 mL was the lowest volume to yield Cq values that did not significantly differ from 25 mL of urine (Tukey-adjusted comparisons; **Figure 1**). As 25 mL was the highest volume of urine investigated in this study, these data suggest that a volume of 5.4 mL of urine is a sufficient volume to assess bacterial load.

**Figure 1.**
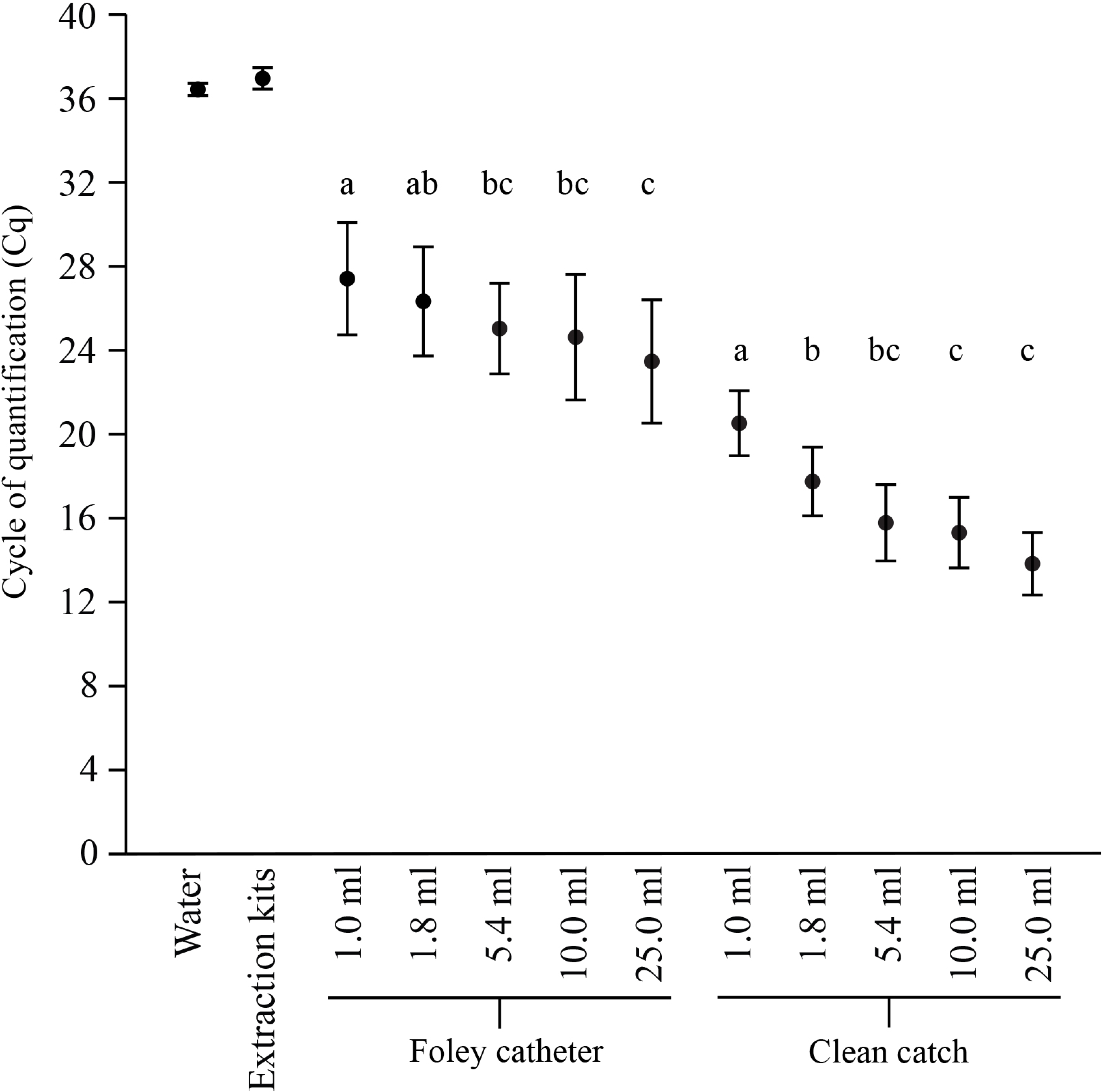
Quantitative real-time PCR (qPCR) of the 16S rRNA gene from urine sample volumes of 1.0, 1.8, 5.4, 10, and 25 ml. Both Foley catheter and clean catch urine collection methods yielded samples with microbial burdens exceeding those of blank DNA extraction kits (Foley catheter, N = 4, clean catch N = 5). Letters correspond to pairwise comparisons where p > 0.05, suggesting the microbial load in those volumes were not different from each other. Plotted values are mean ± standard error.

#### 16S rRNA gene profiles of urine samples

Bacterial profile richness (Chao1 index) and evenness (Shannon and Inverse Simpson indices) were evaluated for differences in alpha diversity between urine samples and negative technical controls and urine samples processed at different volumes. Alpha diversity did not vary between Foley catheter or clean catch urine and negative technical controls, nor did volume of urine processed after correcting for multiple comparisons **(Supplementary Table 2)**. A global effect of sample volume on evenness was observed for Foley catheter urine; however, no pairwise comparisons were significant after correcting for multiple comparisons **(Supplementary Table 2)**. These data indicate that bacterial richness and evenness was similar between urine collection methods, urine volumes, and even with negative technical controls, which emphasizes the importance of including technical controls, as the contaminant DNA sequences as assessed by alpha diversity can be similar to biological samples.

Bacterial profile composition (Jaccard index) and structure (Bray-Curtis index) were evaluated for differences in beta diversity in the bacterial profiles of Foley catheter and clean catch urine samples with negative technical controls. The bacterial profiles of all five volumes of Foley catheter and clean catch urine samples significantly differed from those of negative technical controls (**Table 1**, **Figure 2**). When assessing the influence of urine volume on bacterial profiles, subject identity, not urine sample volume, principally influenced the composition and structure of urine bacterial profiles, regardless of the method of collection (**Table 1**, **Figure 2**). These data indicate the bacterial profiles of urine from either collection method are distinct from technical controls and volumes as low as one milliliter are likely sufficient to characterize the taxa present in the urine of pregnant women.

**Figure 2.**
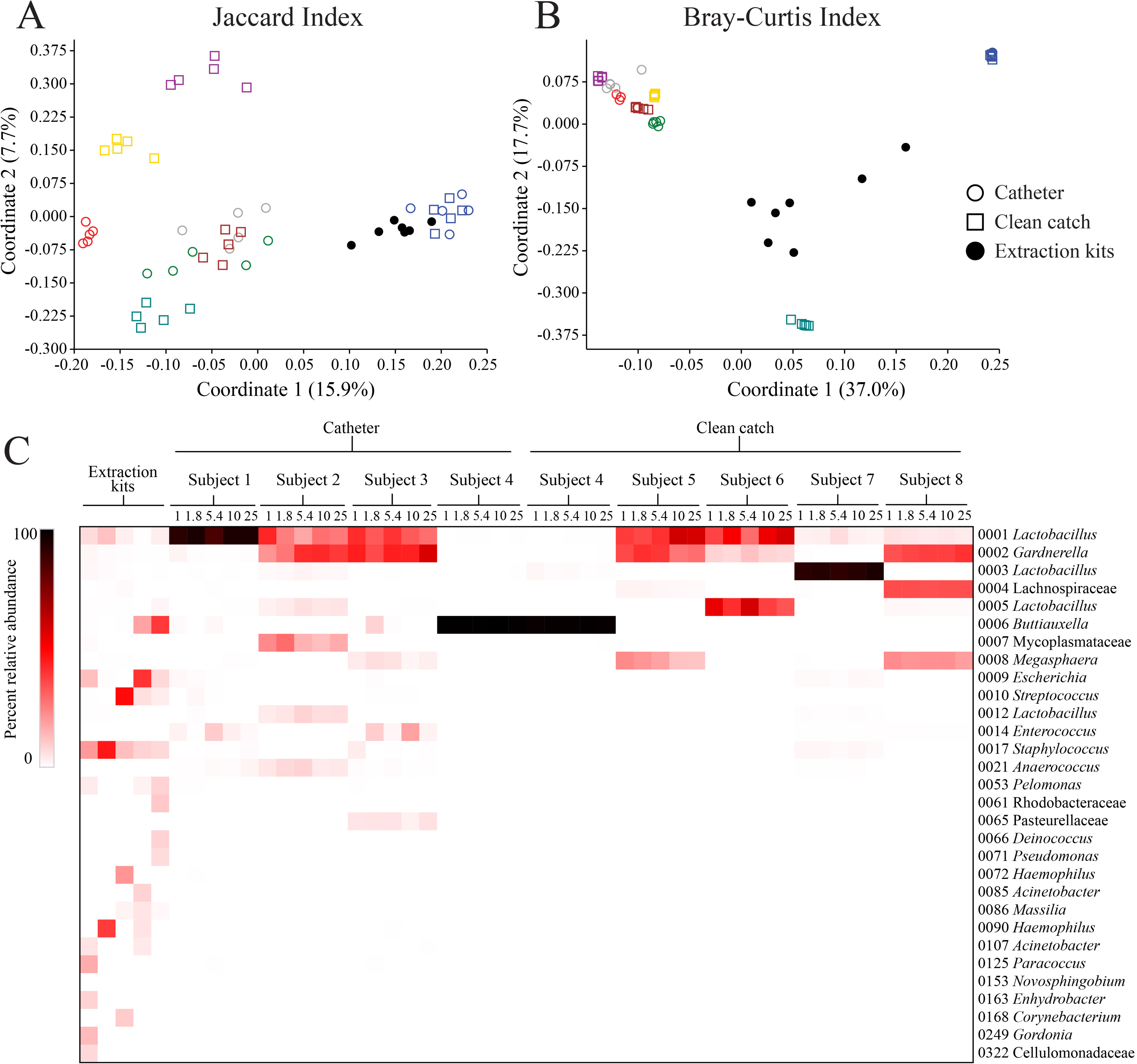
Principal Coordinates Analysis (PCoA) plots and a heatmap illustrating variation in bacterial profiles of Foley catheter urine, clean catch urine, and technical controls. Panels A and B illustrate that the composition and structure of the bacterial profiles of all urine samples, independent of sample volume or collection method, were distinct from those of DNA extraction kit controls. Subject identity, indicated by color, was the principal driver of urine bacterial profiles. In panel C, prominent OTUs (≥1% average relative abundance) among urine samples are visualized from eight pregnant women ordered by urine collection method and sample volume. Urine volume had little influence on overall bacterial profiles, while subject identity was the principal factor. The bacterial profiles of blank DNA extraction kits are distinct from urine and are indicated on the left.

**Table 1.** Statistical analysis of bacterial community composition (Jaccard similarity index) and structure (Bray-Curtis similarity index) for Foley catheter urine and clean catch urine processed at five different volumes and compared to blank controls.

| **Jaccard NPMANOVA** | **F** | **R<sup>2</sup>** | **P** |
| --- | --- | --- | --- |
| **Foley catheter v Blank controls** |  |  |  |
| 1.0 ml | 1.0893 | 0.0901 | **0.0056** |
| 1.8 ml | 1.1143 | 0.0920 | **0.0046** |
| 5.4 ml | 1.1300 | 0.0932 | **0.0019** |
| 10 ml | 1.2040 | 0.0912 | **0.0008** |
| 25 ml | 1.1190 | 0.0923 | **0.0086** |
| **Clean catch v Blank controls** |  |  |  |
| 1.0 ml | 1.1223 | 0.0926 | **0.0026** |
| 1.8 ml | 1.1113 | 0.1000 | **0.0031** |
| 5.4 ml | 1.1021 | 0.0993 | **0.0041** |
| 10 ml | 1.0916 | 0.0903 | **0.0062** |
| 25 ml | 1.0863 | 0.0980 | **0.0095** |
| **Foley catheter** |  |  |  |
| Subject (n = 4) | 1.5412 | 0.2241 | **0.0001** |
| Volume | 1.0023 | 0.1943 | 0.4393 |
| **Clean catch** |  |  |  |
| Subject (n = 5) | 1.7169 | 0.2555 | **0.0001** |
| Volume | 1.0028 | 0.1492 | 0.4313 |
| **Bray-Curtis NPMANOVA** | **F** | **R<sup>2</sup>** | **P** |
| **Foley catheter v Blank controls** |  |  |  |
| 1.0 ml | 2.7119 | 0.1978 | **0.0011** |
| 1.8 ml | 2.5168 | 0.1862 | **0.0011** |
| 5.4 ml | 2.4866 | 0.1844 | **0.0009** |
| 10 ml | 3.4612 | 0.2239 | **0.0008** |
| 25 ml | 2.4820 | 0.1841 | **0.0011** |
| **Clean catch v Blank controls** |  |  |  |
| 1.0 ml | 2.2496 | 0.1698 | **0.0012** |
| 1.8 ml | 1.6062 | 0.1384 | **0.0321** |
| 5.4 ml | 1.6077 | 0.1385 | **0.0299** |
| 10 ml | 2.3175 | 0.1740 | **0.0020** |
| 25 ml | 1.6200 | 0.1394 | **0.0353** |
| **Foley catheter** |  |  |  |
| Subject (n = 4) | 12.5868 | 0.7004 | **0.0001** |
| Volume | 1.0389 | 0.0771 | 0.4032 |
| **Clean catch** |  |  |  |
| Subject (n = 5) | 27.4836 | 0.8427 | **0.0001** |
| Volume | 1.1295 | 0.0346 | 0.3222 |

#### Study Component 1 outcome

As expected, urine volume influenced bacterial load to a point, such that greater volumes yielded a larger bacterial load up to 5.4 mL. Notably, volume did not impact the diversity of urine samples, suggesting volumes as low as 1 mL are sufficient to characterize the bacterial diversity in urine samples. Still, given that a sample volume of 5.4 mL was the lowest volume of urine to yield Cq values that did not differ from those of 25 mL of urine, regardless collection method, a urine sample volume of 5.4 mL was used in Component 2 of the study.

### Study Component 2: Evaluating differences in microbial burden and 16S rRNA gene profiles between Foley catheter and mid-stream clean catch urine in relation to those of vaginal swabs

#### Patient characteristics

Twenty-five women were included in Component 2 of the study. The median and interquartile range for age were 24 (21-29) years, for body mass index were 31.7 (26.3-35.8) kg/m^2^, for gestational age at sampling were 39.3 (39-39.85) weeks, and for neonatal birthweight were 3,165 (2,892.5-3,615) grams **(Supplementary Table 1)**. Twenty-two women were African American, two were Caucasian, and one was self-reported as Other. Seven women (28%) had a history of at least one lifetime UTI, and two (8%) experienced a UTI episode earlier during this pregnancy. However, none had a UTI within 30 days of sampling/delivery (data not shown). Five subjects were in labor at the time of sampling (i.e., cervical dilation of 4 cm or greater and cervical dilation rate of 1 cm per hour or greater). Samples from patients in labor (N = 5) were compared to those not in labor (N = 20) for each sample type (i.e. Foley catheter urine, clean catch urine, vaginal swabs) to assess whether labor influenced bacterial diversity. No difference or variation was observed between the two groups for both alpha and beta diversity metrics (Supplementary Table 2) indicating labor did not influence bacterial diversity in urine and vaginal samples in this study. As a result, labor status was therefore not considered in subsequent analyses.

#### Bacterial cultivation

In total, all clean catch samples (n = 25) were culture positive while only 16 of 25 Foley catheter samples were culture positive. Four types of bacteria (*Lactobacilllus* species, *Staphylococcus* species, *Streptococcus* species, and *Ureaplasma urealyticum*) were each cultured from at least 25% (≥ 13 of 50) of all urine samples. Each was cultured less frequently from urine obtained through a Foley catheter than through mid-stream clean catch urine (**Figure 3**). These results are expected given that these bacterial taxa are common members of the vaginal and skin microbiota, suggesting nearby mucosal sites may be contaminating clean catch urine samples. *Ureaplasma urealyticum* was the most frequently cultured bacteria in Foley catheter urine (13/25 women), suggesting that this genital mycoplasma is a typical resident microbe of the urinary tract. This is significant because *Ureaplasma* species are often considered pathogenic and known to be associated with adverse pregnancy outcomes, however, these women were asymptomatic through delivery.

**Figure 3.**
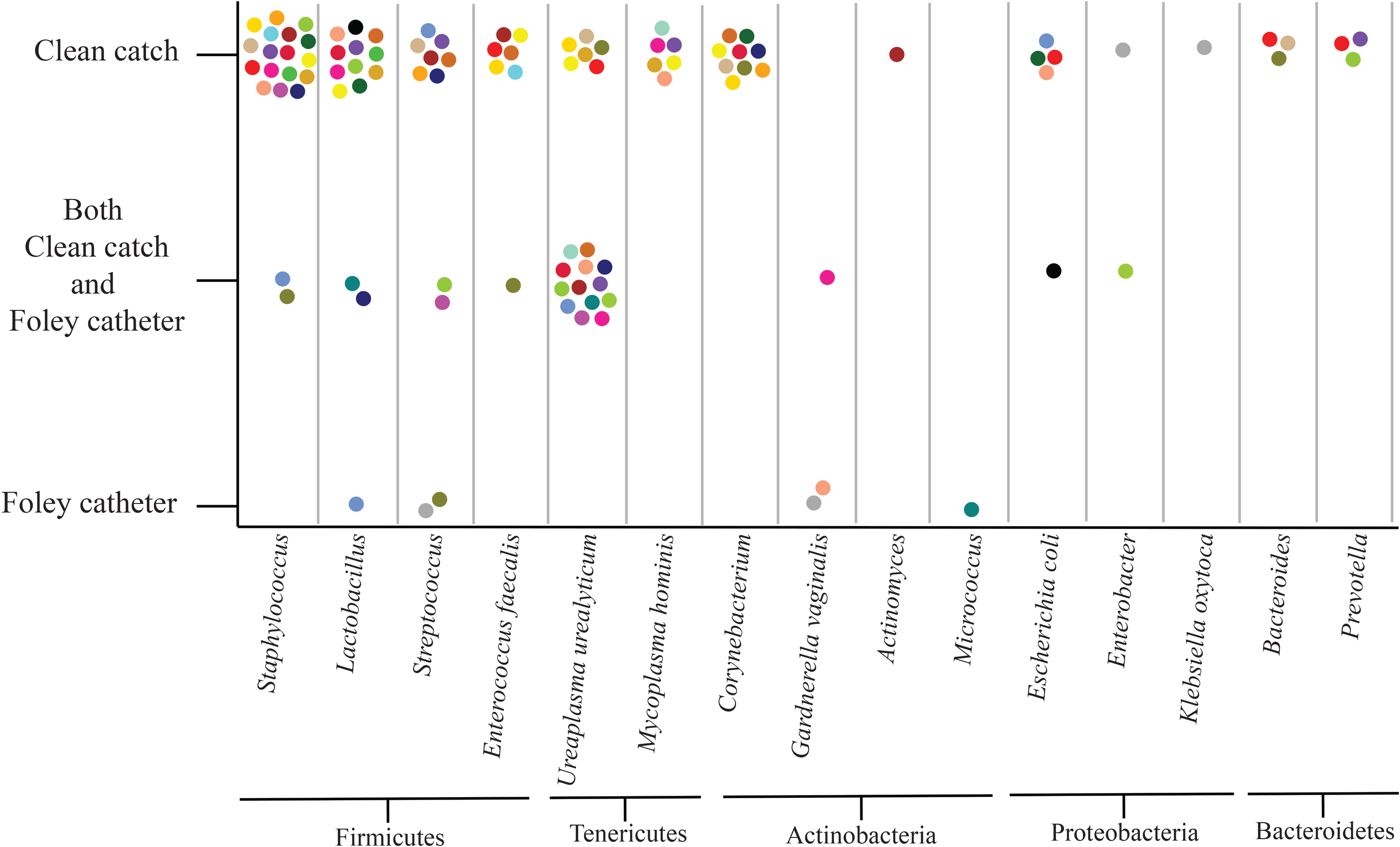
Urine bacterial cultivation results indicating recovery of bacterial phylotypes from catheter urine, clean catch urine, or both. Subject identity is indicated by color. In all but two occurrences, *Staphylococcus* species recovered were coagulase negative, the exception being *S. aureus* recovered in both urine samples of one patient and the clean catch urine of the other patient.

#### Quantitative real-time PCR (qPCR) of 16S rRNA gene abundance in urine and vaginal swab samples

Differences in the bacterial loads of Foley catheter urine, clean catch urine, and vaginal swabs were assessed via qPCR. The bacterial load of clean catch urine (Wilcoxon matched-pairs: W = 325, p < 0.0001)) and vaginal swabs (W = 321, p < 0.0001) exceeded that of catheter urine (**Figure 4A**). The bacterial loads of clean catch urine and vaginal swabs were comparable (W = 210, p = 0.201), suggesting these sample types share similar microbial biomass. While the bacterial load of Foley catheter urine and clean catch urine samples exceeded that of technical controls, Foley catheter urine contained a significantly lower microbial biomass than both clean catch urine and vaginal swab samples. This is important because samples with lower bacterial load have a greater susceptibility to influence from contaminating DNA, which may obscure legitimate members of the microbiota.

**Figure 4.**
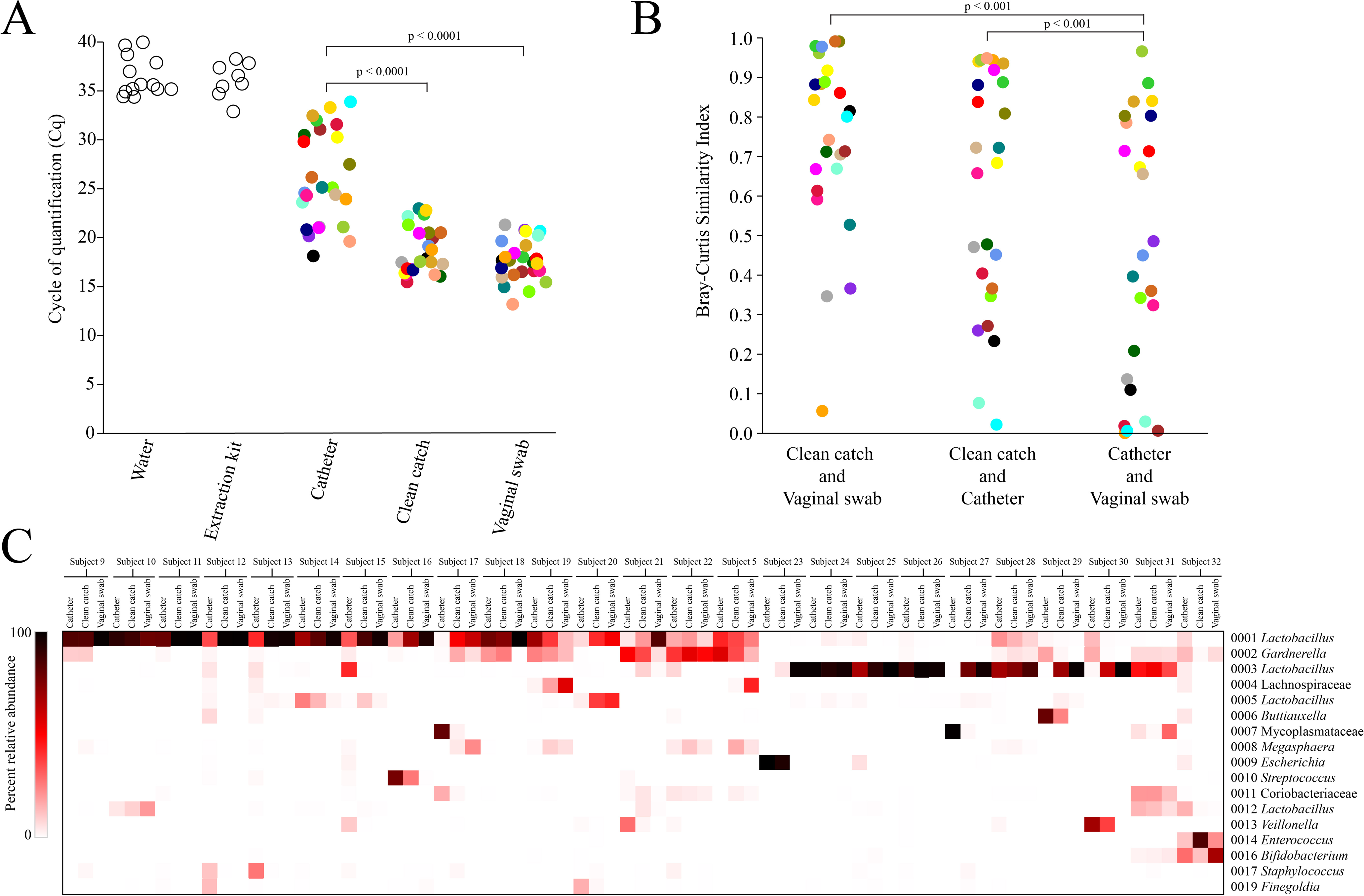
Comparisons of bacterial load and community structure of Foley catheter urine, clean catch urine, and vaginal swabs from 25 pregnant women. In panel A, quantitative real-time PCR (qPCR) of the 16S rRNA gene with cycle of quantification (Cq) values were averaged over multiple runs. Panel B illustrates the similarities in bacterial community structure for each pair of sample types. Panel C includes a heatmap illustrating variation in the profiles of prominent OTUs (≥1% average relative abundance) among paired Foley catheter urine, clean catch urine, and vaginal swab samples from 25 pregnant women. Subjects were ordered via hierarchical clustering of Bray-Curtis similarity indices of clean catch urine samples. Colors indicate subject identity for panels A and B.

#### 16S rRNA gene profiles of paired catheter urine, clean catch urine, and vaginal swab samples

Alpha diversity was assessed for variation between the three sample types (Foley catheter urine, clean catch urine, vaginal swabs). The sample types did not vary in richness (Chao1 index; Friedman’s ANOVA: p = 0.905), but they did vary in heterogeneity based on Shannon (Chi^2^ = 7.28, p = 0.027) and Inverse Simpson (Chi^2^ = 7.44, p = 0.025) indices **(Supplemental Figure 1A & B)**. Specifically, the bacterial profiles of Foley catheter and clean catch urine were not different from each other (Wilcoxon matched pairs with Bonferroni corrections applied: Shannon index: W = 198, p = 1.00, Inverse Simpson index: W = 257, p = 0.900), but both were more heterogeneous than those of vaginal swabs (Foley catheter, Shannon index: W = 266, p = 0.0161, Inverse Simpson index: W = 257, p = 0.033; clean catch, Shannon index: W = 277, p = 0.0062, Inverse Simpson index: W = 253, p = 0.045).

Variation in beta diversity was primarily driven by subject identity for composition and structure of the bacterial profiles of Foley catheter urine, clean catch urine, and vaginal swabs (**Table 2**, **Supplementary Figure 1C & D)**. Nevertheless, controlling for subject identity, the composition of the three sample types, as assessed through the Jaccard index, differed from one another (**Table 2**, **Supplemental Figure 1C)**. With respect to structure, the bacterial profiles of catheter urine, as characterized through the Bray-Curtis index, differed from those of clean catch urine (p = 0.0222) and vaginal swabs (p = 0.0460), but the structure of the bacterial profiles of clean catch urine did not differ from the profiles of their paired vaginal swabs (**Table 2**, **Figure 4B**, **Supplementary Figure 1D)**.

**Table 2.** Statistical analysis of bacterial community composition (Jaccard similarity index) and structure (Bray-Curtis similarity index) for Foley catheter urine, clean catch urine, and vaginal swabs.

| **Jaccard NPMANOVA** | **F** | **R<sup>2</sup>** | **P** |
| --- | --- | --- | --- |
| **Foley Catheter v Clean Catch v Vaginal Swab** |  |  |  |
| Subject (n = 25) | 1.02349 | 0.32957 | **0.0019** |
| Sample Type | 0.98388 | 0.02640 | 0.7941 |
| **Foley Catheter v Clean Catch** |  |  |  |
| Sample Type within subject | 1.0515 | 0.02144 | **0.0022** |
| **Clean Catch v Vaginal Swab** |  |  |  |
| Sample Type within subject | 1.1345 | 0.02309 | **0.0014** |
| **Foley Catheter v Vaginal Swab** |  |  |  |
| Sample Type within subject | 1.1306 | 0.02301 | **0.0027** |
| **Bray-Curtis NPMANOVA** | **F** | **R<sup>2</sup>** | **P** |
| **Foley Catheter v Clean Catch v Vaginal Swab** |  |  |  |
| Subject (n = 25) | 1.55064 | 0.42797 | **0.0002** |
| Sample Type | 0.87084 | 0.02003 | 0.6228 |
| **Foley Catheter v Clean Catch** |  |  |  |
| Sample Type within subject | 0.88327 | 0.01807 | **0.0222** |
| **Clean Catch v Vaginal Swab** |  |  |  |
| Sample Type within subject | 0.92679 | 0.01894 | 0.4834 |
| **Foley Catheter v Vaginal Swab** |  |  |  |
| Sample Type within subject | 1.9757 | 0.03953 | **0.0460** |

The similarity in structure (Bray-Curtis) of the bacterial profiles between clean catch urine and vaginal swabs suggested potential vulvovaginal contamination of the clean catch urine samples. To further investigate this idea, we used the analytical tool SourceTracker to infer the degree to which the bacterial profiles of a source (in this case, vaginal swabs) explained bacterial signals of designated sinks (clean catch or catheter urine). SourceTracker found there was a greater contribution of Operational Taxonomic Units (OTUs) explained by vaginal swabs (source) in clean catch urine (sink) than in catheter urine (sink) (**Supplementary Figure 2**; Wilcoxon paired test: W = 279, p < 0.001). These results suggest the close proximity of the vagina to the urethra increased vulvovaginal contamination of clean catch urine samples; an issue reduced through Foley catheter collection of urine allowing discernment of a distinct urinary microbiota.

The predominant taxa detected in bacterial profiles of catheter urine, clean catch urine, and vaginal swabs were *Lactobacillus* and *Gardnerella* (**Figure 4C**). Basic local alignment search tool (BLAST) analyses indicated that OTUs 1 and 2 were *Lactobacillus iners* and *Gardnerella* spp. (100% identity with type strains *G. leopoldii* UGent06.41 and *G. swidsinskii* GS 9838-1), respectively. OTUs 3, 5, and 12 were each identified as multiple species of *Lactobacillus.* OTU 3 shared identity with *Lactobacillus crispatus*, *L. acidophilus,* and *L. gallinarum* (hereafter considered the more common urogenital lactobacilli, *L. crispatus* [2, 6, 14, 47, 59, 60]), OTU 5 shared identity with *L. jensenii* and *L. fornicalis* (hereafter considered *L jensenii,* the more common urogenital lactobacilli of the two [2, 6, 14, 47, 59, 60]), and OTU 12 shared identity with *L. gasseri* and *L. johnsonii* (hereafter considered *L. gasseri*, the more common urogenital lactobacilli of the two [2, 6, 14, 47, 59, 60]). Thus, consistent with prior observations [6, 13, 60–62], the urogenital bacterial profiles of pregnant women were largely comprised of three community state types: 1) dominance by *Lactobacillus crispatus*; 2) dominance by *Lactobacillus iners*; or, 3) co-dominance by *L. iners* and *Gardnerella* species. Both *Lactobacillus* and *Gardnerella* were rarely detected in the cultivation surveys of urine, which is consistent with the fact that both genera can require specialized media when recovering them from mixed bacterial communities (e.g., *Lactobacillus* De Man, Rogosa, and Sharpe (MRS) agar and V agar, respectively). Besides *Lactobacillus* and *Gardnerella,* the bacterial profiles of catheter urine were variable in high relative abundance taxa including five OTUs with at least one sample with >50% relative abundance: an unclassified Mycoplasmataceae (OTU7), *Escherichia* (OTU 9), *Buttiauxella* (OTU 9)*, Streptococcus* (OTU 10), and *Veillonella* (OTU 13). Notably, five additional OTUs were detected in at least 12 catheter urine samples and had a relative abundance of ≥ 10% in at least one sample. These OTUs included an unclassified Coriobacteriaceae (OTU 11), *Staphylococcus* (OTU 17), *Finegoldia* (OTU 19), *Ureaplasma* (OTU 26), and *Peptoniphilus* (OTU 27).

Linear discriminant analysis effect size (LEfSe) analyses identified eight OTUs that were consistently more abundant in controls than any of the biological sample types, suggesting these OTUs are likely contaminants **(Supplementary Figure 3)**. These OTUs were identified as *Escherichia* (OTU 9), *Staphylococcus* (OTU 17), *Pelomonas* (OTU 53), *Massilia* (OTU 86), *Haemophilus* (OTU 90), *Virgibacillus* (OTU 102), *Acinetobacter* (OTU 107), *Cloacibacterium* (OTU 329). Analyses comparing catheter urine to vaginal swabs identified 17 OTUs more relatively abundant in catheter urine (**Figure 5**). While this analysis did not control for patient identity, four of these seventeen OTUs were also more relatively abundant in catheter urine than negative controls (**Figure 5**). These included *Finegoldia* (OTU 19), *Ureaplasma* (OTU 26), *Anaerococcus* (OTU 49), and an unclassified Clostridiales [OTU 43 (BLAST percent identity to *Fenollaria massiliensis*: 100%)]. The observation that these 4 OTUs were more abundant in catheter urine than in controls or in vaginal swabs suggested they could be potential members of a urinary and bladder microbiota. To further test this idea, their abundances in catheter urine and vaginal swabs were compared directly through paired testing. All four OTUs were more relatively abundant (Wilcoxon matched pairs: OTU 19, W = 152, p = 0.02; OTU 26, W = 134, p = 0.006; OTU 43, W = 69, p = 0.019; OTU 49, W = 74, p = 0.046) and occurred more frequently in catheter urine than vaginal swabs **(Supplementary Figure 4A**). Notably, there were no OTUs that were more enriched in vaginal swabs than they were in Foley catheter urine samples (**Figure 5**). This can be explained because the vaginal swabs were dominated by a few OTUs, accounting for over 90% of the average OTU abundance, and these OTUs were also identified in the urine samples (OTU 1 *Lactobacillus*, OTU 2 *Gardnerella*, OTU 3 *Lactobacillus*, OTU 4 unclassified Lachnospiraceae, OTU 5 *Lactobacillus*, and OTU 16 *Bifidobacterium*).

**Figure 5.**
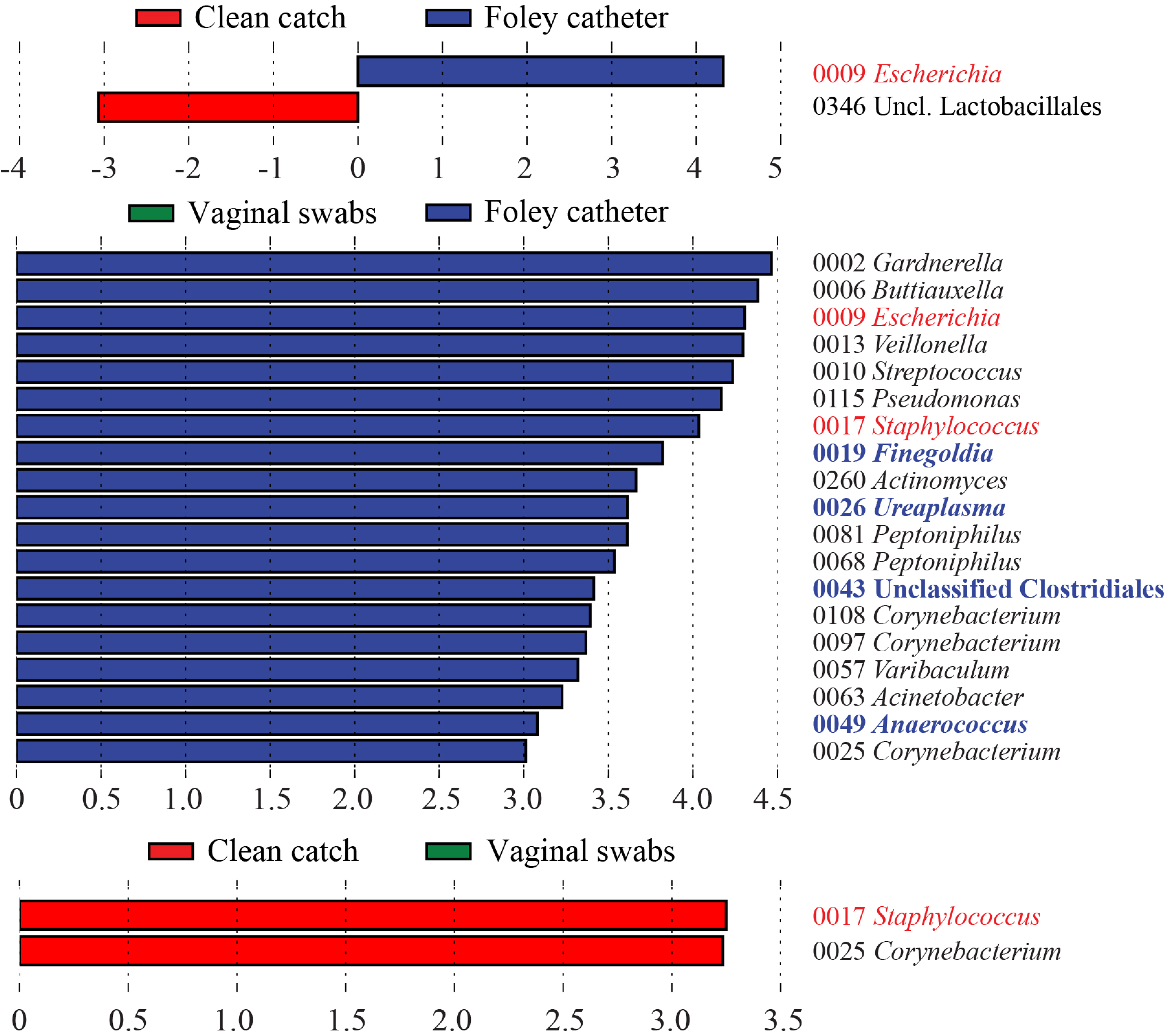
Linear discriminant analysis Effect Size analyses identified several bacteria that were more relatively abundant in Foley catheter urine over vaginal samples, suggesting they are members of a urine and bladder microbiota. Analyses of sequence datasets from Foley catheter urine with clean catch urine, and vaginal swabs. Highlighted in blue are taxa identified as being more relatively abundant in catheter urine than in both clean catch urine and vaginal swabs.

## DISCUSSION

### Principal findings of the study

(1) Quantitative real-time PCR showed that the bacterial load of urine exceeded technical controls regardless of the urine collection method (Foley catheter or clean catch) or the volume of urine processed; (2) A urine volume of 5.4 mL was the lowest to yield a similar 16S rRNA gene load and profile as 25 mL of urine, which was the largest urine volume evaluated; (3) Bacterial isolates were recovered from all 25 clean catch urine samples while only 13 Foley catheter urine samples yielded ioslates; (4) Overall, four culture isolates were detected in at least 25% of all urine samples (*Lactobacilllus* species, *Staphylococcus* species, *Streptococcus* species, and *Ureaplasma urealyticum*) and all four were detected less frequently in catheter than clean catch urine; (5) Molecular sequencing techniques showed that the bacterial profiles of clean catch urine were more similar to vaginal swabs than were catheter urine, suggesting a greater influence of vulvovaginal contamination on clean catch samples; (6) *Lactobacillus iners, L. crispatus,* and *Gardnerella vaginalis* were the most relatively abundant bacteria among all 3 sample types; (7) *Ureaplasma* (detected in culture and molecular surveys) and multiple Gram-positive species including elusive Gram-positive anaerobic cocci (GPAC) such as *Finegoldia* and *Anaerococcus* were more relatively abundant in catheter urine than clean catch urine or vaginal swabs.

Overall, our study suggests that live bacteria commonly exist in the urinary tracts of pregnant women, and that while there is overlap with the vaginal microbiota, there are also distinct lower abundance taxa in samples collected directly from the bladder by catheterization that warrant targeted investigation as potential members of a resident bladder microbiota.

### Urinary Tract Infections (UTIs) and Pregnancy

Approximately 10% of women report having at least one episode of UTI in the previous 12 months and the lifetime probability of a woman having at least one UTI event is around 60% [37, 63–65]. Among women with culture-confirmed UTIs, around 28% had recurrence within six months [37, 66]. The bacterium most responsible for UTIs is *Escherichia coli*, followed by *Staphylococcus saprophyticus*, *Streptococcus agalactiae*, and *Klebsiella* and *Enterococcus* species [22, 23, 39, 44]. The occurrence of a UTI during pregnancy is associated with significant increased odds ratios of 1.4 for low birth weight, 1.3 for preterm birth (< 37 weeks), 1.4 for maternal hypertension and preeclampsia, 1.6 for maternal anemia (hematocrit less than 30%), and 1.4 for chorioamnionitis [23, 24]. The risk of occurrence of these complications is higher among patients with pyelonephritis [23, 67]. Acute pyelonephritis occurs in 2% of pregnant women and recurs in 33% of them [23, 68].

In 2017, a European interdisciplinary group consisting of 17 representatives of 12 medical societies was formed to update the diagnosis and management of uncomplicated UTIs in non-pregnant premenopausal women and concluded that the diagnosis of uncomplicated cystitis be based on clinical criteria evaluating the symptoms of the patient and the course of the disease [69]. They also recommended that no microbiological examination is needed in asymptomatic non-pregnant patients [69]. In pregnancy, ASB is to be detected and treated because of the potential serious complications, including pyelonephritis in 30-40 % of the cases [44]. The American college of Obstetricians and Gynecologists (ACOG) and other societies recommend urine culture as one of the routine tests to be obtained early in pregnancy [70, 71], with mid-stream clean catch being the most common collection approach. If mid-stream clean catch urine culture is performed, the detection of *Escherichia coli* is predictive of bacterial UTI while the detection of other bacteria such as enterococci and group B streptococci is not predictive [69].

### Vulvovaginal contamination of urine samples

The bacterial profiles of all three urogenital sample types from the pregnant women in this study showed a dominance of *Lactobacillus* and *Gardnerella.* One interpretation of these results is that there is a continuum of bacterial sharing between the vagina and the lower urinary tract niches. An alternative explanation is that bacteria present in “clean catch” urine are simply contaminants from the vagina. Mid-stream clean catch remains a more convenient approach for patients and health care staff; however, culture results of clean catch samples need careful interpretation [72–74]. A study of 113 asymptomatic pregnant women showed a high level of contamination in clean catch samples [72]. Baerheim et al. [73] found that employing precautions such as cleaning the perineum or obtaining mid-stream samples led to similar contamination rates as obtaining samples without any precautions [73]. Lifshitz et al. [74] evaluated 242 symptomatic female patients divided into 3 groups 1) no cleaning, 2) perineal cleaning and midstream sampling, and 3) perineal cleaning, midstream sampling and vaginal tampon. Contamination rates in the three groups were all similar at approximately 30% [74]. In women undergoing cesarean deliveries, two randomized clinical trials showed an increase in the incidence of UTIs in the indwelling bladder catheterization group compared to the no catheterization group [75–77]. Mid-stream clean catch thus remains a more convenient approach for patients and health care staff; however, culture results of clean catch samples need careful interpretation [72–74]. Our cultivation studies confirmed that Foley catheter samples yield less cultivars than clean catch samples and are less likely to be contaminated by vaginal microbiota than are clean catch samples.

### Support for the existence of a urinary bladder microbiota during pregnancy

For Study Component 2, while individual identity was the primary influence on 16S rRNA gene profiles, differences were found for both beta diversity indices (i.e. Jaccard and Bray Curtis) between the three sample types: catheter urine, clean catch urine, and vaginal swabs. Pairwise comparisons showed that the bacterial profiles of catheter urine were different than those of both clean catch urine and vaginal swabs in composition and structure, whereas the profiles of clean catch urine were different from vaginal swabs only in structure. This suggests that urine collected via the clean catch method has a bacterial profile more similar to that of the vagina due to proximity and increased likelihood of vulvovaginal contamination. In this way, clean catch urine likely represents a mix of members of the skin and vaginal microbial communities in addition to the microbiota of the urinary bladder. Whilst urine collected via a catheter is still likely subject to some degree of vulvovaginal and skin contamination, our data support the current consensus in the field that catheterization limits contamination and that bacteria present in catheterized urine be referred to as the urinary bladder microbiota (4).

LEfSe analyses identified seven OTUs that were more relatively abundant in Foley catheter urine than in blank extraction controls, and four of these OTUs were also more relatively abundant in catheter urine than in vaginal swab samples, suggesting that these OTUs may be representative of members of the bladder microbiota, at least in some women. While only *Ureaplasma* was recovered from cultivation surveys, *Finegoldia* and *Anaerococcus* species are Gram-positive anaerobic cocci recalcitrant to culture, typically requiring long incubation times and complex growth requirements [78]. This may explain why they were not recovered in culture. Other groups investigating urine via molecular surveys have also identified *Finegoldia* and *Anaerococcus* species in non-pregnant females, in healthy cohorts as well as among women with non-UTI urinary disorders [14, 50–52, 59, 79]. The fourth OTU, while originally described as an unclassified Clostridiales, was a 100% sequence match to the newly identified genus and species *Fenollaria massiliensis* gen. nov., sp. nov [80, 81]. Its relatively recent discovery may explain why it was not detected by culture as most clinical microbiology laboratories are likely unfamiliar with this organism. This microorganism is discussed in more detail below.

*Ureaplasma* species, like other members of the class Mollicutes, lack a cell wall, require specialized media, and can require long incubation times [43, 82]. *Ureaplasma* is well-documented in its association with diseases of the urinary tract [4, 53, 59, 79, 82, 83] as well as adverse pregnancy outcomes, including preterm birth [84–89] and diseases of the neonate [90–93]. In fact, it is the most common microorganism found in the amniotic cavity [88, 94]. Despite its associations with disease, *Ureaplasma* is frequently detected by culture and molecular surveys in urine of asymptomatic women, pregnant [35, 41, 43, 47, 48, 95] and non-pregnant [50, 59, 60, 79], suggesting its potential role in a bladder microbiota. The various consequences associated with the presence of *Ureaplasma* in the bladder of pregnant women is likely multifactorial, of which adverse outcomes are likely associated with a combination of the individual’s own immune response [96], the composition and structure of the broader microbial community [11, 97], and the specific strain of *Ureaplasma* [98–101].

*Finegoldia* has been associated with the genitourinary tract, gastrointestinal tract, and skin as a commensal but has also been isolated from and attributed to infections from wounds and various body sites making it an opportunistic pathogen. Difficulty in cultivation has been evident in clinical reports where accurate diagnoses were dependent on detection via PCR despite cultures yielding negative results [102–104].

*Anaerococcus* species are also commensals of the skin, gastrointestinal tract, and oral cavity and members have been isolated from vaginal secretions and purulent wounds [78, 105, 106]. Literature reports successful growth of *Anaerococcus* species on standard anaerobic plate types [107, 108] by some groups, while others indicate the addition of supplemental nutrients such as hemin and vitamin K [78, 107–110]. Reported incubation times have also varied from 2 days up to 7 days [108, 110].

*Fellonaria massiliensis* is a newly discovered and understudied anaerobic rod recovered and characterized from osteoarticular, genital, and tissue samples, and is suggested to be a genital-associated microbe [80, 81]. The two studies describing this organism report growth on several enriched media types after 72 hours and on supplemented Brucella Blood Agar after 48 hours under anaerobic conditions.

Many of the less abundant genera detected in recent studies represent species that are not captured by routine cultures or have never been successfully cultured. Thus, it is also important to perform viability assays, such as expanded quantitative urine culture (EQUC) methods, to demonstrate viability of bacteria from these samples because molecular surveys do not differentiate between ubiquitous environmental 16S rRNA gene sequences and those from living bacteria [2]. Therefore, the studies above demonstrate that to elucidate the existence and the characteristics of urine microbiota it is important to include large sample sizes, rigorous and extensive culture techniques, and ample technical controls to account for background DNA contamination.

Our study showed that catheter urine samples do yield a 16S rRNA gene signal beyond that evident in controls and suggest catheterization may be an appropriate sampling method for evaluating any microbial communities that may exist in the bladder. Also, our results suggest that the vaginal microbiota influences or contaminates clean catch urine to a larger degree than catheter urine, and that while some catheter samples are still influenced by vaginal microbes, a potentially unique signal may exist in some individuals. Our evidence suggests that underlying *Lactobacillus* and *Gardnerella* abundance in the urine of pregnant women, anaerobic organisms like *Finegoldia,*

*Anaerococcus*, and *Peptoniphilus* may be low abundant members of a bladder microbial community. However, to more confidently assert that the bladder contains microbiota in pregnancy, a suprapubic sampling approach would provide better insight. We suspect that the gold standard for investigating a potential bladder microbiota would be suprapubic aspiration of urine in concert with tissue sampling of the bladder epithelium. Culture should be performed to discriminate live bacteria from remnant DNA from dead bacteria.

The results reported herein are clinically relevant given that they support the fact that a positive culture during routine urine analyses in asymptomatic women with <100,000 CFU may indicate the detection of components of the normal urine microbiota rather than a specific pathogen, and thus antibiotic treatment is not required. Moreover, our results highlight the importance of the method of urine collection, as well as its volume when interpreting microbiological results in the urine during pregnancy, emphasizing the possible contamination of urine with vaginal bacteria when collected with clean catch method, which is the classic method of collection.

### Strengths and limitations

This is the first extensive study that attempts to characterize the urinary microbiota in pregnancy by comparing Foley catheter, clean catch, and vaginal samples from 25 women. In addition, this is the first study that compared different volumes of urine to determine the optimal volume for performing 16S rRNA gene surveys. Furthermore, this study utilized cultivation, qPCR, and sequencing approaches to study the existence and viability of microbiota in the bladder. The main limitation of the study is that our population mainly consists of one ethnic group (i.e. African American). It is possible that other ethnic groups may have a different bladder microbiota. Non-pregnant women were not included in this study, therefore differences and similarities between a healthy female bladder microbiota and a pregnant female bladder microbiota cannot be addressed. Additionally, it is difficult to assess if and how much of the bacterial signal in catheter urine was due to vulvovaginal contamination, specifically regarding the top three most abundant taxa. More extensive culture methods may have allowed the lower abundance anaerobic organisms to be recovered.

### Conclusions

Our study suggests that resident bacterial communities exist in the bladder and urine, and that there is overlap with the vaginal microbiota. While the most frequent microorganisms recovered by Foley catheter samples were *Ureaplasma,* molecular surveys identified low abundance anaerobic bacteria in addition to *Ureaplasma* as potential members of a bladder microbiota.

### Future directions

Future research should endeavor to evaluate the typical presence of a microbial community within the urinary tract of pregnant women by comparing and analyzing 16S rRNA gene sequence data from pregnant women delivering preterm (condition; defined as delivering ≤ 37 weeks) and at term (biological control; defined as delivering > 37 weeks) and the appropriate technical controls. In doing so, we can assess influences of resident microbiota on perinatal health and pregnancy outcomes and identify bacteria or bacterial communities whose presence or absence can serve as potential risk indicators. Ultimately, being able to categorize and describe bladder microbial communities and their associations with preterm birth should lead to potential targets, therapeutic interventions, and other methods for treating or modifying the bladder microbiota and lessening the impact of urinary tract-associated pregnancy complications.

## MATERIALS AND METHODS

### Study subjects and sample collection

#### Study subjects

This was a cross-sectional study in which the urinary and vaginal microbiota were examined in 25 women admitted for delivery to the Detroit Medical Center (Detroit, USA) over a four-month period. All participants provided written informed consent for the collection and use of samples for research purposes under the protocols approved by the institutional review boards of Wayne State University and the *Eunice Kennedy Shriver* National Institute of Child Health and Human Development, National Institutes of Health, Department of Health and Human Services. Inclusion criteria were: 1) delivery after 36 weeks of gestation, and 2) intact membranes at the time of collection of vaginal swabs and clean catch urine samples. Exclusion criteria were: 1) any maternal or fetal condition that requires termination of pregnancy; 2) known major fetal anomaly or fetal demise; 3) active vaginal bleeding; 4) serious medical illness (e.g. renal insufficiency, congestive heart disease, chronic respiratory insufficiency, etc.); 5) asthma requiring systemic steroids; 6) patient requiring anti-platelet or non-steroidal anti-inflammatory drugs; 7) active hepatitis; and 8) signs or symptoms of asymptomatic bacteriuria (ASB), urinary tract infection (UTI), and/or pyelonephritis at the time of sampling. In this study, UTI was defined as >100 CFU of a single bacterial type per milliliter (CFU/mL) of urine [111] coincident with one of the following symptoms: hematuria, dysuria frequency, urgency or suprapubic pressure [36, 37]. ASB was defined as the presence of 100,000 CFU/mL without any of the above-mentioned UTI-associated symptoms [35–37]. Pyelonephritis was indicated by the presence of systemic signs or symptoms such as fever, nausea and vomiting, chills or flank pain [35–37].

In the first component of the study, which assessed bacterial load and profiles of urine samples at multiple volumes compared to background contamination, no subject had received antibiotics in the last week. In the second component of the study, which entailed evaluating differences in bacterial load and profiles of Foley catheter urine and clean catch urine and vaginal swabs, no subject had received antibiotics in the last month.

#### Sample collection

On admission, each woman provided a mid-stream clean catch urine specimen (CC). Women were instructed to begin micturition for 3 seconds, stop, cleanse the area with an iodine-based wipe, and provide a clean-catch, midstream urinary specimen in a sterile cup. A speculum exam was then performed, and a sample of vaginal fluid was collected from the posterior vaginal fornix under direct visualization by an obstetrician using a FLOQSwab (Copan Diagnostics, Murrieta, CA, USA). A sterile Foley catheter was inserted during labor or prior to a cesarean delivery (8.36 ± 1.93 (mean ± SE) hours after the original clean catch sample was collected), and a second urine specimen was collected through the catheter. Urine (excluding the aliquots used for culture, see below) and vaginal swabs were frozen at -80°C within one hour of collection.

#### Bacterial culture of urine

Two milliliter aliquots of urine were sent for bacterial culture to the University Laboratories Microbiology Core in the Detroit Medical Center, wherein they were processed and cultured under aerobic and anaerobic conditions on the same day of sample collection. A genital mycoplasma assay (Mycofast US; Logan, UT) was also conducted for each urine sample [112]. In each case, one drop of urine, equivalent to approximately 0.05 mL, was used. Incubation for aerobic, anaerobic and *Mycoplasma*/*Ureaplasma* cultures was performed at 35°C. TSA 5% SB (Trypticase Soy Agar w/5% Sheep’s Blood), Columbia CNA SB, MacConkey and MTM II (Modified Thayer Martin) were incubated aerobically with 8% CO_2_. Brucella OxyPRAS Plus, KVL/BBE Biplate (Brucella Laked Blood Agar with Kanamycin and Vancomycin/Bacteroides Bile Esculin Agar) and CDC ANA BLD (CDC Anaerobic Blood Agar) were incubated anaerobically in a non-CO_2_ incubator. The genital mycoplasma assays were performed under an oxic environment without CO_2_ supplementation. Urine cultures were incubated for 48 hours. The taxonomies of resultant isolates were characterized using Matrix-Assisted Laser Desorption/Ionization Time-Of-Flight Mass Spectrometry (MALDI-TOF/MS) within the University Laboratories Microbiology Core [113].

#### Genomic DNA extractions

##### Preparation of urine samples for DNA extraction

For 1, 1.8, and 5.4 mL sample volumes, urine samples were originally stored at -80°C in either 2 mL cryovials or in 15 mL centrifuge tubes. Samples were thawed at room temperature and thoroughly vortexed before aliquoting into 1.8 mL microcentrifuge tubes. Samples were spun in a microcentrifuge in a 4° C cold room for 30 minutes at 17,000*g*. After centrifugation, each sample had the majority of supernatant removed. For the 1 mL sample, approximately 750 µl of supernatant was carefully removed with a 1 mL pipette tip, avoiding the pellet, thereby leaving about 250 µl of the supernatant and the pellet for DNA extraction. For the 1.8 mL sample, 775 µl was removed twice, again being careful to avoid disturbing the pellet, leaving about 250 µl of the supernatant and the pellet for DNA extraction. For the three 1.8 mL tubes constituting the 5.4 mL sample, 860 µl was removed twice from each tube, carefully avoiding the pellet, leaving 80 µl of supernatant and the pellet in each tube for DNA extraction. The initial step of the DNA extraction protocol requires adding 500 µl of the kit’s PowerBead Solution to the sample; the PowerBead Solution was added directly to these 1.8 mL tubes. The tubes were then thoroughly mixed through vortexing and by pipetting the solution up and down to ensure that the pellet was dislodged into solution and would be transferred to the bead tube in the next extraction step. For the special case of 5.4 mL samples which were split into three tubes, 500 µl of PowerBead Solution was added to the first tube and the mixture was transferred to the subsequent tubes for further mixing until the contents of all tubes were thoroughly mixed together and then transferred to PowerBead Tubes for DNA extractions.

For DNA extractions performed on 10 and 25 mL samples, samples were thawed at room temperature and thoroughly vortexed before transferring 10 or 25 mL into 50 mL centrifuge tubes. These samples were spun at 4°C at 17,000*g* for 30 minutes. After centrifugation, the supernatant was removed without disturbing the pellet leaving approximately ≤ 500 mL. The initial step of the DNA extraction protocol requires adding 500 µl of PowerBead Solution to the sample, so the PowerBead Solution was added directly to these 50mL tubes. These tubes were then thoroughly mixed through vortexing and pipetting the sample up and down to ensure that the pellet was dislodged into solution and would be transferred to the bead tube for the following step in the extraction protocol.

##### Extraction protocol

Genomic DNA was extracted from urine and vaginal swab samples using QIAGEN DNeasy PowerLyzer PowerSoil Kits according to the manufacturer’s protocol with the following modifications to increase DNA yield of low microbial biomass samples: 1) instead of adding 750 μl of PowerBead Solution to each sample, 500 μl of PowerBead Solution and 200 μl of phenol/chloroform:isoamyl alcohol were added and the sample was incubated in the PowerBead Tubes at room temperature for 10 minutes, 2) steps that entail adding Solutions C2 and C3 were combined into one step; 1 μl of RNase A enzyme was also added, 3) instead of adding 1200 μl of Solution C4, 650 μl of C4 and 650 μl of 100% ethanol were added, 4) the dry-spin after Solution C5 was extended from 1 to 2 minutes, 5) Solution C6 was heated to 60°C prior to elution of DNA, and 6) 60 μl instead of 100 μl of Solution C6 were added to the Spin Column and incubated for 5 minutes before final centrifugation. Blank DNA extraction kits with no urine sample added (n = 12) were processed alongside urine samples. Purified DNA was stored at -20°C.

#### Quantitative real-time PCR (qPCR) of 16S rRNA genes in samples

Bacterial DNA load within samples was determined via quantitative real-time PCR (qPCR) amplification of the V1 – V2 region of the 16S rRNA gene according to a protocol described by Dickson et al. [114] with minor modifications. These modifications included the use of a degenerative forward primer (27f-CM: 5’-AGA GTT TGA TCM TGG CTC AG-3’) and a degenerate probe containing locked nucleic acids (+) (BSR65/17: 5’-56FAM-TAA +YA+C ATG +CA+A GT+C GA-BHQ1-3’). All qPCR reactions were performed in triplicate (20 μl each), with each reaction containing 0.6 μM of 27f-CM primer, 0.6 μM of 357R primer (5’-CTG CTG CCT YCC GTA G-3’), 0.25 μM of BSR65/17 probe, 10.0 μl of 2X TaqMan Environmental Master Mix 2.0 (Life Technologies, Carlsbad, CA), and 4.0 μl purified DNA. Cycling conditions were as follows: 95°C for 10 min, and 45 cycles of 94°C for 30 s, 50°C for 30 s, and 72°C for 30 s. Fluorescent readings were taken at the end of each cycle on an ABI 7500 Real-Time PCR System (Applied Biosystems, Waltham, MA). Raw amplification data were normalized to the ROX passive reference dye and analyzed with Standard Curve 3.3.0-SR2-build15 (Thermo Fisher Cloud), using automatic threshold and baseline settings. Cycle of quantification (Cq) values were calculated and defined as the average number of cycles required for normalized fluorescence to exponentially increase. DNA derived from *Escherichia coli* ATCC 25922 containing seven 16S rRNA gene copies per genome (GenBank accession: CP009072) was quantified using a Qubit 3.0 fluorometer with a Qubit dsDNA HS Assay kit (Life Technologies, Carlsbad, CA) according to the manufacturer’s instructions and used for the generation of standard curves. To estimate qPCR efficiency, a standard curve containing seven 10-fold serial dilutions (three replicates each) ranging from 1.99 X 10^7^ to 1.99 X 10^1^ copies was included in each run. Prior to analyzing qPCR data with the on-line platform Thermo Fisher Cloud (Standard Curve (SR) 3.3.0-SR2-build15), an external master standard curve was generated by performing a regression of the standard curve data from all six qPCR runs (slope = -3.4629, y-intercept = 40.122, R^2^ = 0.9798).

#### 16S rRNA gene sequencing

The V4 region of the 16S gene was amplified using a modified PCR approach (95° for 2 min, followed by 32 cycles of 95° for 30 s, 55° for 30 s, and 72° for 30 s, with a final elongation step at 72° for 10 min). DNA template volumes were 5 μl for urine and blank DNA extraction kits, and 3 μl for vaginal swabs. 16S rRNA gene sequencing was completed on an Illumina MiSeq (San Diego, CA) instrument at the University of Michigan’s Center for Microbial Systems (Ann Arbor, MI) using the dual indexing sequencing MiSeq protocol strategy developed by Schloss and colleagues [115, 116].

#### 16S rRNA gene sequence processing

Sequence data were processed using Mothur software (v1.39.5) [117]. Specifically, paired reads were assembled, quality-filtered (no ambiguous base calls, homopolymers ≤ 8 bases long), and aligned to the SILVA 16S rDNA reference database (release 102) [118, 119]. Sequences in the final dataset had an average length of 253 bp. We performed a preclustering step (diffs = 2) to reduce potential influence of sequencing errors and removed chimeras identified by UCHIME [120]. For taxonomic classification, the SILVA reference database [119] was used with a confidence threshold of 80% [121]. Sequences from an unknown domain, Eukaryota, Chloroplasts, Mitochondria, or Archaea were removed. Operational taxonomic units (OTUs) were defined using a 3% sequence dissimilarity cutoff. Good’s coverage values for all urine and vaginal samples exceeded 99%.

#### Statistical analyses

##### Bacterial culture

The rate of cultivation of bacterial phylotypes (as identified via MALDI-TOF, e.g. *Lactobacilllus*, coagulase negative *Staphylococcus*, *Ureaplasma urealyticum*) from urine was compared between Foley catheter and clean catch collection methods using generalized estimating equation models assuming a binomial distribution (i.e. detected or non-detected). Only bacterial phylotypes detected in at least 20% of the samples, regardless of the collection method, were tested. Significance of the odds ratios was assessed via Wald tests, implemented in the *geepack* package in R (v 3.4) [122]. The paired differences in the total numbers of bacterial phylotypes detected within the Foley catheter and clean catch urine samples among the women were assessed using a Poisson generalized estimating equation model.

##### 16S rRNA gene qPCR

To assess differences in bacterial load between each urine volume and collection method and blank DNA extraction kit controls, differences in cycle of quantification (Cq) were evaluated via Mann-Whitney tests. To assess variation in 16S rDNA abundance among urine samples of different volumes from the same women, variation in Cq values was evaluated via repeated-measures analysis of variance (ANOVA) followed by Tukey’s tests for pair-wise comparisons or Friedman’s ANOVA followed by Wilcoxon matched pairs tests. In component 2 of the study, differences in 16S rDNA abundance between sample types were assessed using Friedman’s ANOVA followed by Wilcoxon matched pairs tests. Statistical analyses were performed using PAST software (v3.16) [123].

##### 16S rRNA gene profile alpha diversity

For Study Component 1, blank DNA extraction kit controls were sequenced twice and subsequently pooled bioinformatically. The controls with Good’s coverage values exceeding 98% were retained for analysis [n = 5, additional controls were processed during DNA extractions for Study Component 2 and used as part of the LEfSe analysis (n = 7)]. Alpha diversity in Study Component 1 was analyzed after subsampling individual libraries to 447 sequences, which corresponds to the sequencing depth of the least represented background technical control sample. After subsampling, Good’s coverage remained above 95% for all but one sample (91%).

Alpha diversity in Study Component 2 was analyzed after subsampling individual libraries to 2007 sequences, which only excluded one sample. In Component 2, after subsampling, Good’s coverage values for urine and vaginal samples remained greater than or equal to 98%.

Alpha diversity was assessed using the Chao1 index as an indicator of richness and the Shannon and Inverse Simpson indices as indicators of heterogeneity (evenness). Differences in alpha diversity between urine and background technical control samples were evaluated through t-tests or Mann-Whitney tests. For comparisons among different urine volumes or between sample types, variation in alpha diversity was evaluated through repeated measures ANOVA followed by Tukey’s matched-pairs or their non-parametric equivalents. Alpha diversity indices were generated in mothur (v1.39.5) and statistically evaluated in PAST (v3.16).

##### 16S rRNA gene profile beta diversity

To evaluate differences in beta diversity of 16S rRNA gene profiles, Jaccard (i.e. composition) and Bray-Curtis (i.e. structure) similarity index values were calculated using OTU percent relative abundance data within samples. These data were visualized through Principal Coordinates Analyses (PCoA). Non-parametric MANOVA (NPMANOVA) tests were performed on Jaccard and Bray-Curtis similarity indices to assess differences between background technical controls and different urine volumes (Component 1), and variation among catheter urine, clean catch urine, and vaginal samples (Component 2). Beta diversity indices and PCoA plots were generated using PAST software (v3.16). Non-parametric MANOVA [124–126] tests were performed in R (version 3.4.2) with the adonis function in the vegan package. The “strata” parameter in the adonis function was used to control for repeated measures. Heatmaps were generated via the Morpheus online tool [127].

##### SourceTracker analysis

SourceTracker software [128] was used to identify what percentage of OTUs found in urine samples could be attributed to contamination from vaginal samples. For each urine collection method, SourceTracker analysis was done in triplicate with a rarefaction depth of 500 and the proportions from the three model runs were averaged to give the mean percentage of OTUs predicted to be from vaginal samples. Singletons and doubletons (OTUs detected only once or twice) were removed from 16S rRNA gene datasets prior to these analyses. Wilcoxon paired tests of the averaged SourceTracker runs were evaluated in PAST (v3.16).

##### LEfSe analysis

Linear discriminant analysis effect size, or LEfSe, was used to identify OTUs that differed in relative abundance between each of the three biological sample types (Foley catheter urine, clean catch urine, vaginal swabs) and blank DNA extraction kits (n = 7). Singleton OTUs were removed from the datasets prior to analyses and the LDA cutoff score was set to 3.0.

---

**Supplementary Figure 1.**
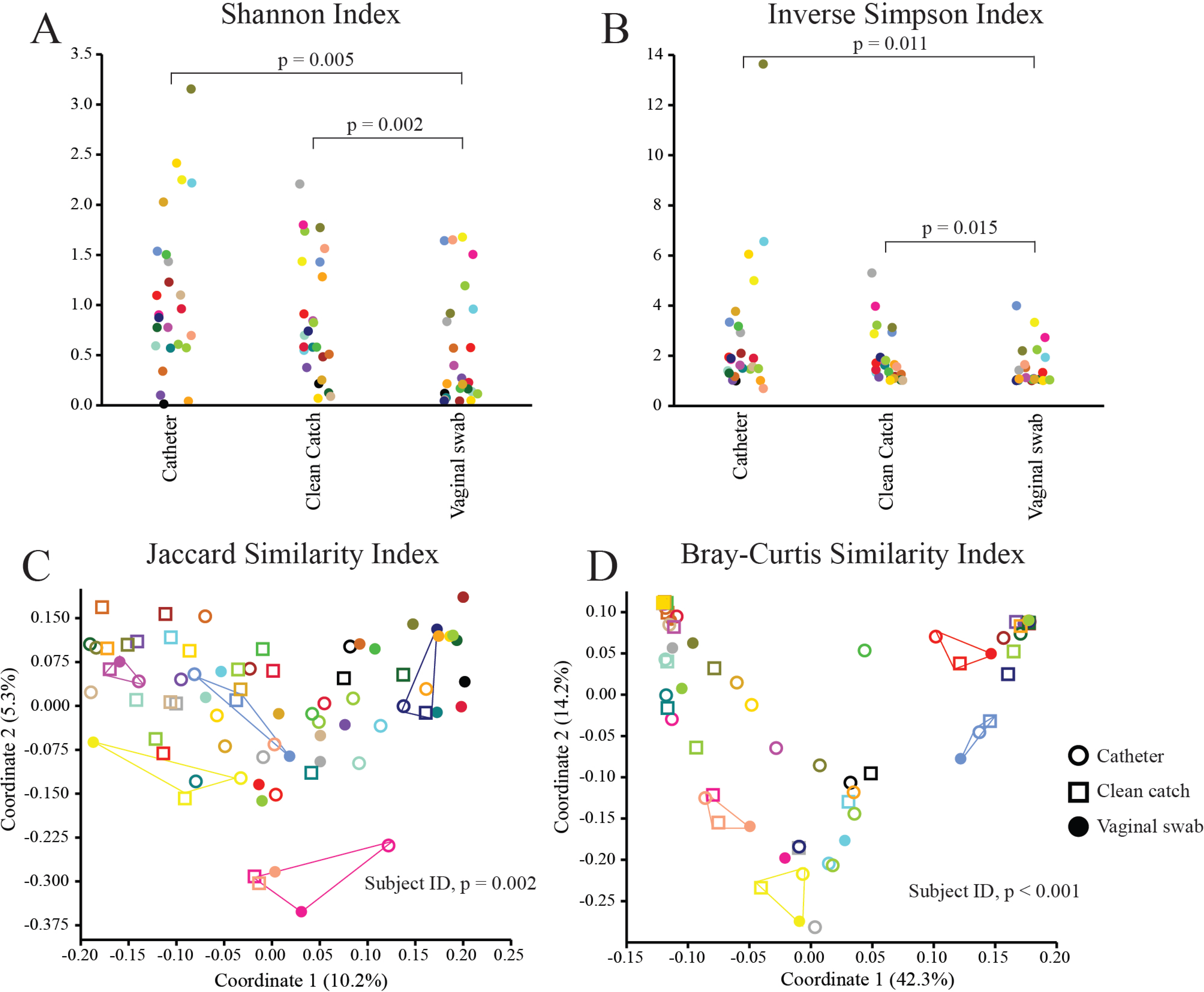
Jitter and Principal Coordinates Analysis (PCoA) plots illustrating alpha and beta diversities of Foley catheter urine, clean catch urine, and vaginal swabs collected from the same women. Panels A and B show differences in heterogeneity between sample types, with catheter urine having the greatest diversity for both indices. Panels C and D illustrate the composition and structure of the bacterial profiles of the three sample types. Several subjects are highlighted to illustrate the influence of individual identity on the bacterial profiles. Subject identity is indicated by the same color scheme across the 4 panels.

**Supplementary Figure 2.**
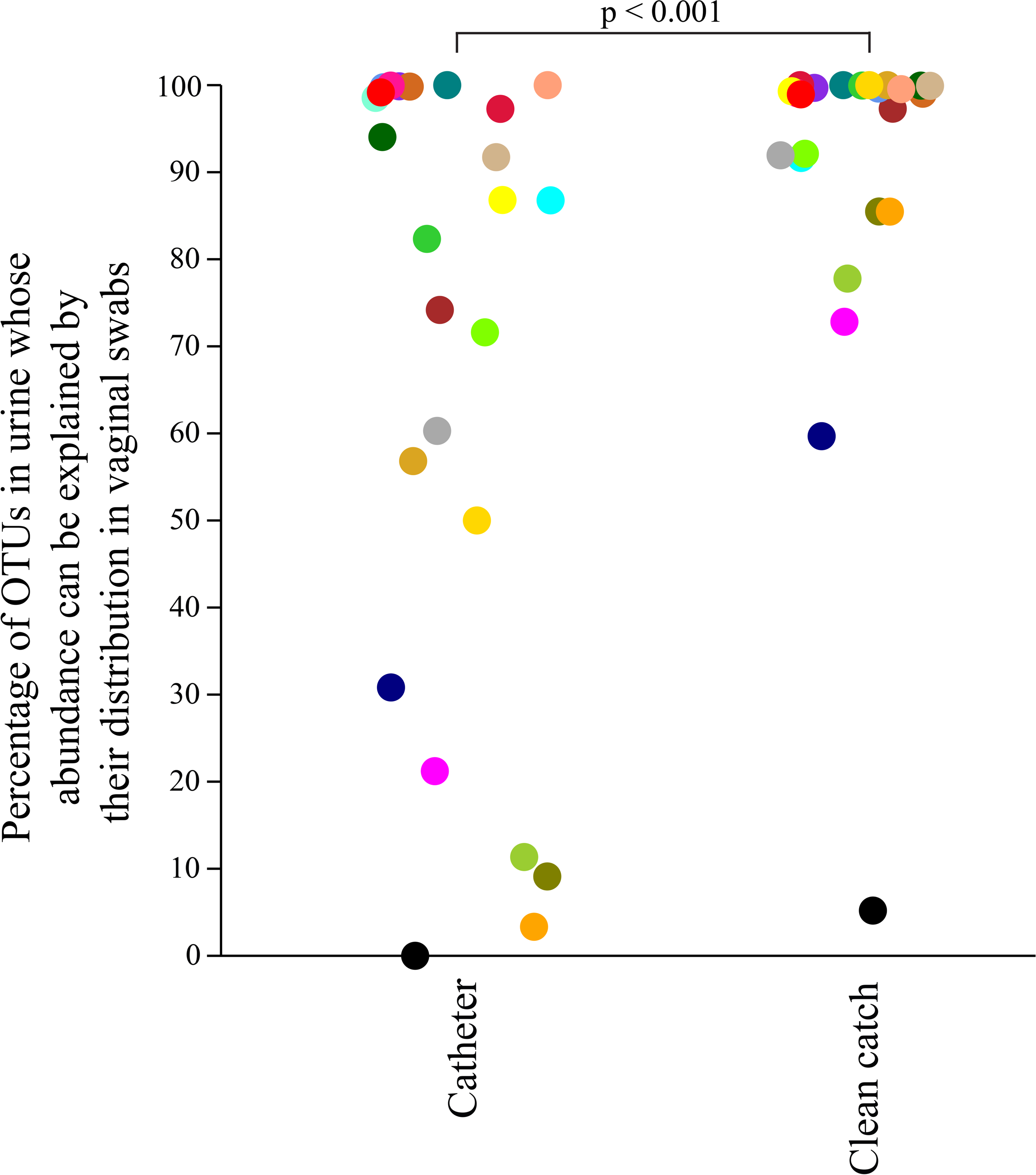
SourceTracker analysis comparing the percentage of OTUs explained by vaginal swabs among Foley catheter urine and clean catch urine. Colors represent subject identity.

**Supplementary Figure 3.**
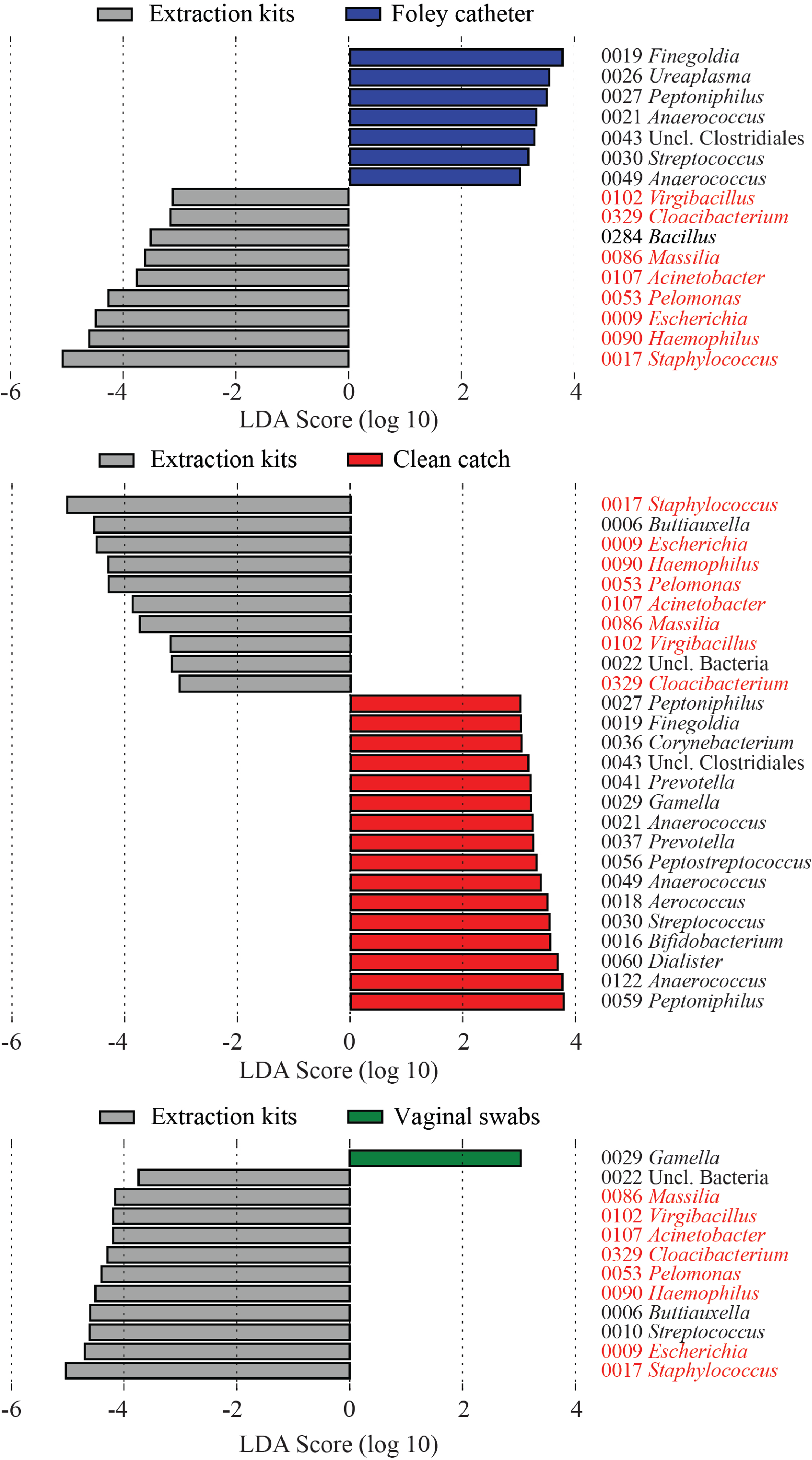
Linear discriminant analysis Effect Size analyses identified several bacteria that were more relatively abundant in blank extraction kits. Analyses of 16S rRNA gene sequence datasets from DNA extraction compared to each sample type. OTUs highlighted in red were more relatively abundant in extractions kits than in all 3 biological samples suggesting they are likely contaminant sequences.

**Supplementary Figure 4.**
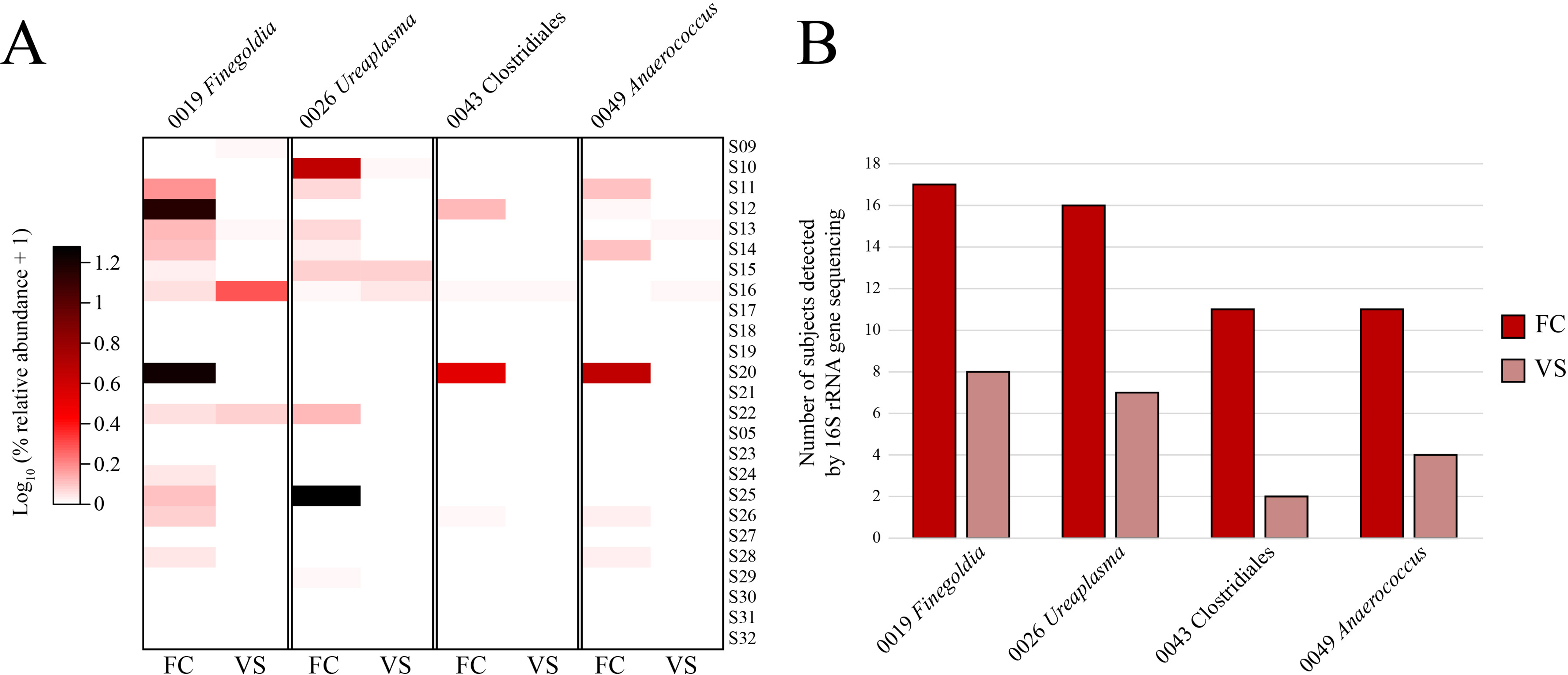
Comparisons of relative abundances and frequency of four OTUs between Foley catheter urine and vaginal swab samples. Each OTU was identified as preferentially abundant in Foley catheter urine samples by LEfSe analysis. The heatmap in panel A includes log transformed relative abundance data for Foley catheter urine (FC) and vaginal swabs (VS). Each row represents an individual and subjects are ordered the same as Figure 4C. Panel B indicates the number of subjects in which each OTU was detected for each sample type.

**Supplementary Table 1.** Descriptive and clinical characteristics of subjects for Study Components 1 and 2.

| Study Component 1 |  |  |
| --- | --- | --- |
| N = 8 | Median | IQR<sup>a</sup> |
| Age (yrs) | 27.5 | 25.2 – 28.2 |
| BMI<sup>b</sup> (kg/m<sup>2</sup>) | 31.6 | 28.5 – 45.8 |
| GA<sup>c</sup> at sampling (wks) | 39.7 | 38.8 – 40.8 |
| Birthweight (g) | 3392 | 3256 – 3882 |
| Race |  |  |
| African American | 8 (100 %) |  |

| Study Component 2 |  |  |
| --- | --- | --- |
| N = 25 | Median | IQR<sup>a</sup> |
| Age (yrs) | 24.0 | 21.0 – 29.0 |
| BMI<sup>b</sup> (kg/m<sup>2</sup>) | 31.7 | 26.3 – 35.8 |
| GA<sup>c</sup> at sampling (wks) | 39.3 | 39.0 – 39.85 |
| Birthweight (g) | 3165 | 2892.5 – 3615 |
| Race |  |  |
| African American | 22 (88.0 %) |  |
| White | 2 (8.0 %) |  |
| Other | 1 (4.0 %) |  |

<sup>a</sup> Interquartile range
<sup>b</sup> Body Mass Index
<sup>c</sup> Gestational Age

**Supplementary Table 2.** Comparisons of alpha diversity of Foley catheter urine and clean catch urine processed at 5 different volumes and blank controls.

| **t-test or Mann-Whitney U test** | **Chao1** |  | **Shannon** |  | **Inverse Simpson** |  |
| --- | --- | --- | --- | --- | --- | --- |
| **Foley catheter v Blank controls** | **t** | **P** | **t** | **P** | **t** | **P** |
| 1.0 ml | 0.2860 | 0.7814 | 1.6081 | 0.1423 | 1.5260 | 0.1613 |
| 1.8 ml | 1.4256 | 0.1877 | 1.3154 | 0.2209 | 1.1052 | 0.2978 |
| 5.4 ml | 2.0131 | 0.0746 | 14* | 0.9247 | 0.8111 | 0.4382 |
| 10 ml | 2.4313 | 0.0379 | 1.9453 | 0.0836 | 1.6474 | 0.1339 |
| 25 ml | 1.4779 | 0.1736 | 1.4726 | 0.1750 | 1.5136 | 0.1644 |
| **Clean catch v Blank controls** | **t** | **P** | **t** | **P** | **t** | **P** |
| 1.0 ml | 0.4407 | 0.6688 | 2.1093 | 0.0611 | 1.7277 | 0.1147 |
| 1.8 ml | 0.0219 | 0.9830 | 2.2362 | 0.0493 | 1.7596 | 0.1090 |
| 5.4 ml | 0.7251 | 0.4850 | 2.3172 | 0.0430 | 1.8545 | 0.0934 |
| 10 ml | 0.2532 | 0.8053 | 2.5737 | 0.0277 | 2.0930 | 0.0628 |
| 25 ml | 0.0377 | 0.9707 | 2.8564 | 0.0171 | 2.1924 | 0.0531 |
| **Repeated-measures ANOVA**<br>**or Friedman's ANOVA** | **F** | **P** | **F** | **P** | **F** | **P** |
| **Foley catheter** |  |  |  |  |  |  |
| Volume | 1.7700 | 0.1998 | 10.45** | **0.0349** | 4.215 | **0.0233** |
| Pairwise comparisons |  |  |  | All > 0.01 |  | All > 0.01 |
| **Clean catch** | **F** | **P** | **F** | **P** | **F** | **P** |
| Volume | 0.5478 | 0.7033 | 1.462 | 0.2600 | 1.911 | 0.1578 |
| Pairwise comparisons |  |  |  |  |  |  |

\*Mann-Whitney U; \*\*chi<sup>2</sup>

